# Modular, part-based control of gene expression response time using protein degradation tags

**DOI:** 10.1101/482331

**Authors:** Ethan M Jones, Callan E Monette, John P Marken, Sejal Dhawan, Theresa Gibney, Christine Li, Wukun Liu, Alyssa Luz-Ricca, Xida Ren, Xingyu Zheng, Margaret Saha

**Author notes:** These authors contributed equally to the work (Ethan M Jones, Callan E Monette).

## Abstract

A fundamental principle of cellular signal processing is the encoding of information within the temporal dynamics of regulatory circuits. If synthetic circuits are to achieve the versatility and effectiveness of naturally-occurring circuits, it is necessary to develop simple, effective methods for the control of the dynamical properties of genetic circuits. However, current approaches to dynamical control often require extensive rewiring of circuit architecture, which hinders their implementation in a variety of systems. Therefore, it is essential that simple, modular, genetic parts-based frameworks are created to control the dynamical properties of circuits. Here we address this need by implementing a modular, genetic parts-based system which tunes the response time of a gene’s expression by tuning its degradation rate via the application of protein degradation tags with various affinities to their protease. This system provides a simple, easily-applicable framework for controlling the temporal aspects of genetic circuit behavior.

## Introduction

Synthetic biology aims to gain predictable control over genetic circuit behavior using modular genetic parts. Currently, synthetic biologists can use modular genetic parts to exert a high level of control over the steady-state behavior of genetic circuits [1,2]. However, as synthetic biologists create more complex genetic circuits designed to mimic or interface with dynamic natural systems, there is a need for frameworks capable of controlling the dynamical properties of genetic circuits. These frameworks should be both simple and modular to encourage their application in a wide variety of systems—ideally, one should be able to adopt such a framework simply by swapping a single genetic part in their constructs.

One fundamental dynamical property of genetic circuits is their response time. In the context of genetic circuits, response time is typically defined as the amount of time for the concentration of a gene’s output to reach half of its steady-state concentration [3]. Since response time is a metric for the amount of time needed for a given circuit to switch from an inactive state to an activated state, control over this property is vital for the design of efficient genetic circuits. It is therefore essential to obtain control over circuit response time using the same modular, parts-based paradigm that is currently used to control the static properties of genetic circuits.

Control over the response time of genetic circuits has previously been described by investigators who utilized post-translational modification [4] or circuit rewiring to include additional feedback loops [5,6]. While these methods indeed offer control over dynamical properties of gene expression, ideally a circuit’s response time could be controlled by changing a single genetic part, much in the same way that a circuit’s steady-state output can be controlled by choice of Ribosome Binding Site. Using the insights from a previously-described theoretical relationship between a gene product’s degradation rate and its response time [3], here we provide a part-based framework for modular control of response time via protein degradation tags of various affinity.

## Results/Discussion

To determine a general solution to the control of genetic circuit response time, we utilized a simple mathematical model of inducible gene expression described by Alon [3]. In this model a gene produces an output protein (x) with a net production rate (α) and a net degradation rate (γ) when expression is induced. This forms the simple ordinary differential equation:

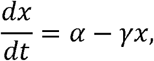

Which can be solved to obtain the concentration of the protein x as a function of time

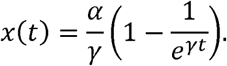

With this equation the steady state concentration of x can be determined to be 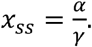 Circuit response time, τ, can then be defined as the amount of time for the concentration of protein x to reach half the steady state value, and the following equation can be obtained:

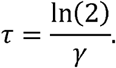

This expression reveals that an inducible gene’s response time is controlled only by the degradation rate of the gene product [3]. Therefore, a modular part-based approach to control a gene’s degradation rate is sufficient to control its response time.

To create a framework for controlling speed via degradation rate, we utilized the *Mesoplasma florum* Lon (mf-Lon) protease system characterized by Gur and Sauer in 2008 [7] which Collins and Cameron developed in 2014 into a set of modular genetic parts [8]. In this system mf-Lon protease degrades proteins which contain a C terminal 27 amino acid protein degradation tag (pdt) in a manner analogous to the degradation of SsrA tags by ClpXP protease in *Escherichia coli* (*E. coli*). Importantly, the mf-Lon protease system developed by Collins and Cameron consists of multiple variant pdts, each possessing a distinct affinity for mf-Lon and hence a distinct degradation rate. We reasoned that this system would provide an accessible and tunable way to control circuit response time via protein degradation rate (Figure 1a).

**Figure 1:**
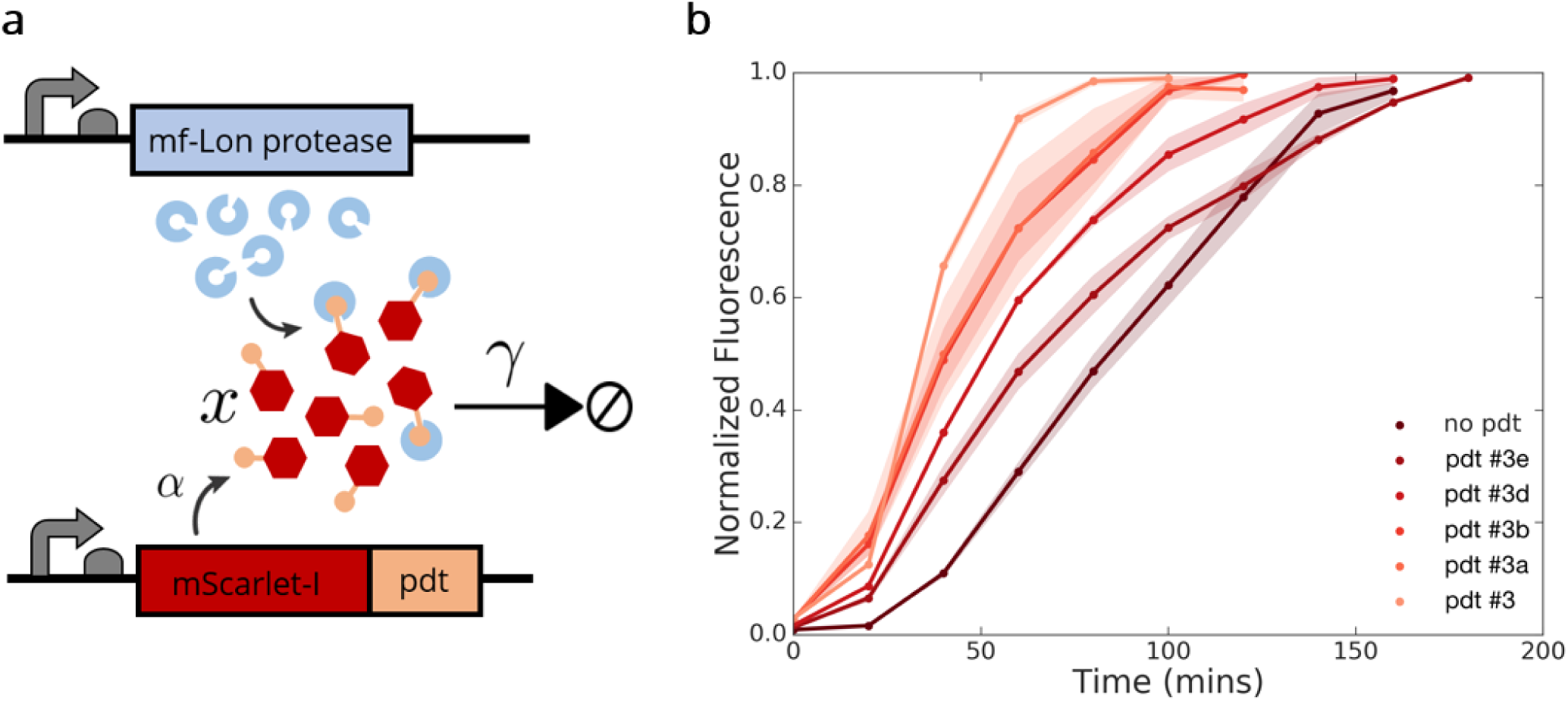
(A) Schematic of response time control system. An inducible fluorescent reporter x with a pdt is produced at rate α and is degraded by the E. coli orthogonal protease mf-Lon, increasing degradation rate (γ) and decreasing response time in a tag-specific fashion. **(B)** Absolute fluorescence measurements in Molecules of Equivalent Fluorescein (MEFL) of gene expression of ATc inducible mScarlet-I + pdt constructs for pdt #3e through #3 from [8], listed in order of increasing protease affinity. These measurements are normalized to their steady state value. The timecourses are truncated when they reach steady state for clearer comparison of this property across conditions (Supplemental Methods). Each data point represents the geometric mean of 10,000 or more single cell measurements taken via flow cytometry of three biological replicates. Shaded region represents one geometric standard deviation above and below the mean.

To confirm that circuit response time could be controlled via degradation rate alone, we created a series of anhydrotetracycline (ATc) inducible constructs expressing pdt tagged fluorescent reporters. We transformed each construct into *E. coli* along with a mf-Lon expressing plasmid and determined circuit response time to an input of ATc by performing time course fluorescence measurements using flow cytometry. We found that the stronger pdts caused constructs to reach steady state faster in response to a fixed ATc input, indicating that degradation rate can indeed be used to tune circuit response time (Figure 1b).

Next, we sought to determine empirically the mathematical relationship between reporter degradation rate and circuit response time. We obtained the relative degradation rates of each construct by comparing their steady state fluorescence relative to an untagged control and found that the relationship between the degradation rates of each pdt were consistent with those reported by Collins and Cameron (Figure 2a) [8]. We then compared the relationship between these degradation rates and circuit response time and found that the two values displayed an inverse relationship, consistent with Alon’s model (Figure 2b).

**Figure 2:**
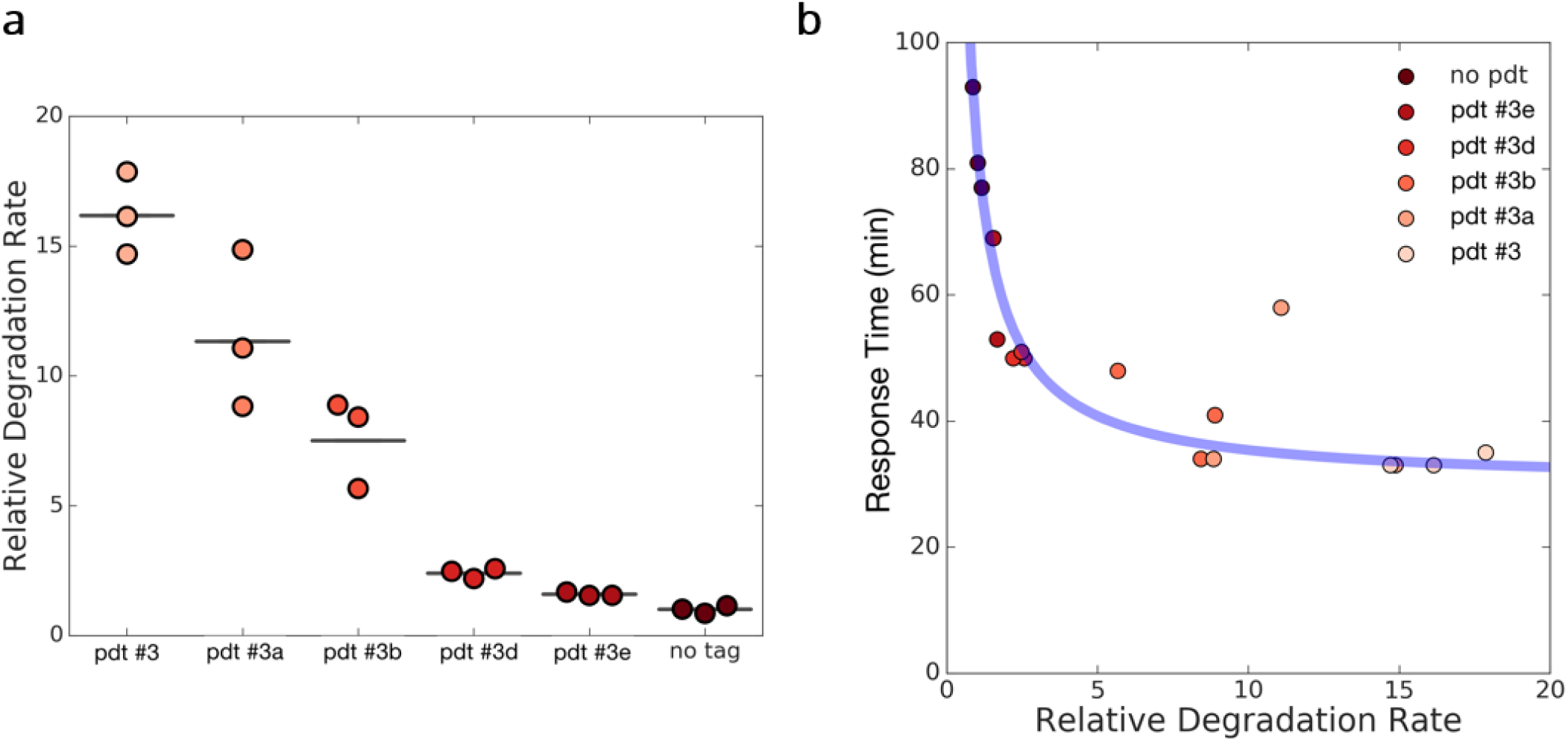
(A) Relative degradation rates measured in ATc inducible mScarlet-I pdt constructs. Each data point represents the population geometric mean of at least 10,000 cells of a distinct biological replicate; the line represents the geometric mean of replicates. **(B)** Comparison of response times shown in Figure 1 versus the relative degradation rates from (A). The blue line is a guide to the eye that demonstrates the inverse scaling between the two variables (Supplemental Methods).

This degradation-based system provides a simple, plug-and-play method for tuning genetic circuit response time, enabling control of the dynamics of synthetic circuits with modular genetic parts. Furthermore, although increasing a gene product’s degradation rate causes a decrease in its steady-state concentration, this decrease can be ameliorated by increasing the production rate of the gene product, as predicted by Alon’s model (Supplemental Figure 1). The consistency displayed between our system and its associated mathematical model ensures a predictable framework for tunable response time and represents a validation of Alon’s model of gene expression dynamics. Our work provides a simple, reliable system for parts-based control over response time, which will grant more precise control over output dynamics and improve the informed design of dynamically encoded circuits.

## Supporting information

## Supporting Information

<u>Abbreviations:</u>

mf-Lon: *Mesoplasma florum* Lon

pdt: protein degradation tag

ATc: anhydrotetracycline

Complete materials and methods, supporting figure S1, supporting table T1 containing information on all constructs used in this study; Supporting sequences, and raw MEFL data used in analysis.

## Author Contributions

EMJ designed constructs and experiments, SD, TVG, EMJ, CEM, CHL, ALR and XZ performed experiments and cloned constructs, EMJ JPM, WL and XR analyzed data, EMJ and CEM wrote manuscript, EMJ, JPM, CEM and MSS edited manuscript, all authors helped conceive project and reviewed and approved manuscript

## Acknowledgements

We would like to thank Andrew D Halleran for his insightful discussions and Dr. Greg Conradi Smith for his assistance with mathematical modeling.

