## Supplementary material for "Modular, part-based control of gene expression response time using protein degradation tags"

**Supplementary Methods**

**Construct Design and Assembly:**

Amino acid sequences for pdts and mf-Lon were obtained from Cameron and Collins 2014, manually codon optimized for *E. coli* using Integrated DNA Technology’s (IDT) codon optimization tool, and then synthesized as gene blocks. The same process was repeated for the fast folding red fluorescent protein mScarlet-I [1]. Gene blocks were then cloned into the backbone pSB1C3. ATc inducible reporter constructs were cloned by using multipart Gibson Assembly to combine the pTet promoter (Bba_R0051), mScarlet-I and the various protein degradation tags (pdt) along with a constitutive TetR construct onto a pSB1C3 backbone. IPTG inducible mf-Lon constructs were cloned in a similar fashion, combining the pLac0-1 (Bba_R0011) promoter with mf-Lon along with a constitutive LacI construct onto the medium copy backbone pSB3K3.

**Circuit response time characterization and readjustment characterization:**

ATc inducible reporter constructs were co-transformed with an IPTG inducible mf-Lon construct in the 10-Beta *E. coli* strain (New England Biolabs). Colonies were picked into M9 minimal media with 0.4% glycerol and grown overnight. In the morning, cultures were 1:100 diluted into fresh media and grown for 4-7 hours. Cell density was quantified and cells were diluted to an optical density (OD) of 0.01 into fresh media containing 0.1mM IPTG and 50ng/mL ATc. Time points were taken from samples every 20 minutes onto ice and then immediately thereafter, at least 10,000 (typically 20,000) single cell measurements were obtained using flow cytometry (FL3 channel Bio-Rad S3e Cell Sorter). Additionally, 10,000 measurements of 8 peak Rainbow Calibration Particles (Spherotek) were collected on each day where measurement occurred. Samples fluorescence was converted into MEFLs using the python package flowcal [2] and the associated calibration beads. Data was analyzed and Figure 1b displays the inducible gene expression profile of a pulsatile IFFL with the mf-Lon protease serving as an inhibitor and the inducers representing the input. For readjustment characterization, procedure was repeated while increasing the concentration of ATc to 100ng/mL.

**Determination of steady state and calculation of τ and relative degradation rate:**

Each timecourse was defined as reaching steady state at the first timepoint for which the two successive measurements were not above the value at that timepoint. The steady-state value for each gene expression timecourse was defined as the fluorescence value obtained at the point the timecourse reached steady state. The measured timepoints were smoothed via a spline fit in Microsoft Excel to determine the response times, τ, for each replicate by interpolation. Degradation rates for a given pdt were calculated as the ratio between the average steady-state fluorescence of the with-pdt conditions (across replicates) and the average steady-state fluorescence of the without-pdt conditions (across replicates) for that tag. In Figure 2b, the guide to the eye is a function of the form

$$\tau=\frac{A}{B\gamma}+C,$$

where τ and γ are the response time and relative degradation rate, respectively, and A, B, C are arbitrary constants.

**Supporting Table T1:**

| **Construct Name (ID)** | **Plasmid Backbone** |
| --- | --- |
| pTet mScarlet-I no tag (wm17_408.gb) | pSB1C3 |
| pTet mScarlet-I pdt#3 (wm17_409.gb) | pSB1C3 |
| pTet mScarlet-I pdt#3a (wm17_410.gb) | pSB1C3 |
| pTet mScarlet-I pdt#3b (wm17_411.gb) | pSB1C3 |
| pTet mScarlet-I pdt#3d (wm17_413.gb) | pSB1C3 |
| pTet mScarlet-I pdt#3e (wm17_414.gb) | pSB1C3 |
| pLac0-1 mf-Lon (wm17_393.gb) | pSB3K3 |

**Supplementary Figures:**

***Supplementary Figure 1:*** *Absolute (MEFL) (left) and steady-state normalized (right) values of mScarlet-I pdt constructs in multiple ATc induction conditions. As predicted by the model, increasing the production parameter (via increasing [ATc]) has minimal effect on the response time of the circuit (right), however increased [ATc] can restore the steady state value to the same value as seen in the no pdt (no degradation) condition.*

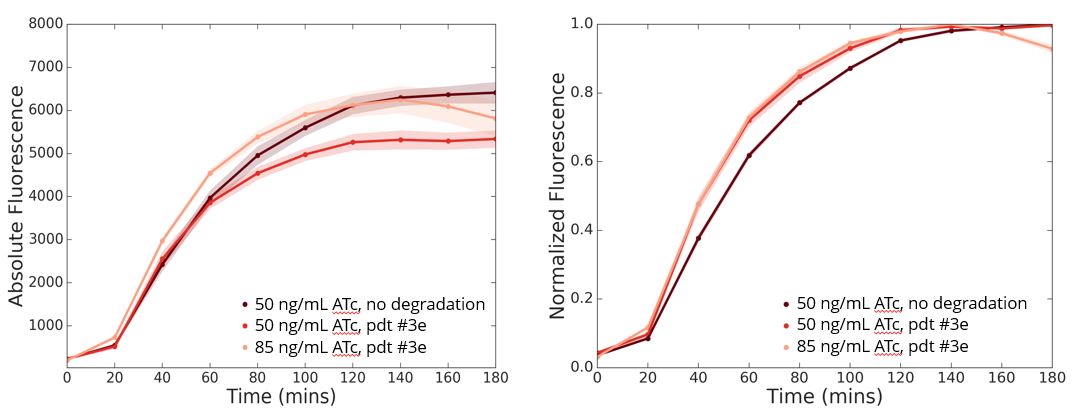
